# Impact of Premorbid Infection on Onset and Disease Activity of Rheumatoid Arthritis

**DOI:** 10.1101/358853

**Authors:** Ruijun Zhang, Jing Li, Jiali Chen, Xiaomei Chen, Xue Li, Chun Li, Yuan Jia, Yunshan Zhou, Limin Ren, Lijun Wu, Jing He, Zhanguo Li

## Abstract

**Objective:** Infections have been implicated in rheumatoid arthritis (RA) development. However, the impact of premorbid infection on initiation and perpetuation of RA has not been well elucidated. Thus, we sought to conduct a large scale on-site survey to study whether premorbid infection may trigger RA and influence status of the disease.

**Methods:** Premorbid infectious events were collected in cohort of 902 RA patients from December 2015 to June 2016. Type of infections prior to RA onset and its possible effects on disease status were analyzed.

**Result:** Three hundred and thirty-four out of 902 patients (37.03%) experienced infections within one month preceding RA onset. The most frequent infections were respiratory (16.08%), intestinal (11.09%) and urinary tract (9.87%) infection, respectively. The infection was associated with increased disease activity. Early onset was found in patients with urinary infection. High disease activity risk was increased in patients who pre-exposure to urinary infection (OR=3.813, 95%CI=1.717-12.418) and upper respiratory infection (OR=2.475, 95%CI= 0.971-6.312).

**Conclusion:** Pre-exposure infections are associated with development of RA. Severe disease status of RA and persistent of active disease status are related to preceding infections.

## Introduction

Rheumatoid arthritis (RA) is a common autoimmune disease characterized by joint destruction and auto-antibodies production.[1] Many studies have demonstrated that infectious agents may contribute to the initiation or perpetuation of RA through a variety of mechanisms. Infection can cause a local inflammatory response. The innate immune system could also be affected by infections agents and then cause RA onset, for instance, pathogen-associated molecular pattern receptors, especially the Toll-like receptors (TLRs) could release inflammatory mediators rapidly after recognizing some preserved structures in bacteria and other infectious agents [2].

Although a definite causative link between a specific infectious agent and the disease has not been established, several arguments support such a possibility. First, in the absence of a certain pathogen, the spectrum of microorganisms involved in triggering RA may include poly-microbial communities or the cumulative effect of bacterial or virus factors [3]. Secondly, infections didn’t lead to RA in all cases, but initiate it in a certain subset of patients who was born with a genetic susceptibility [4-7]. Thirdly, some arthritis occurred based on pre-exposure to microorganism. Several animal models of arthritis are dependent on TLR2, TLR3, TLR4 or TLR9, for instance, rodents injected with streptococcal cell walls (TLR2 ligand) develop severe polyarticular arthritis and TLR4 ligand also play a role in passive K/BxN arthritis [8]. Many studies have shown that components derived from infectious agents can cause autoimmune reaction by molecular mimicry and other mechanisms. Epstein-Barr virus (EBV) is a polyclonal B lymphocyte activator which can increase the production of RF [4]. Oral pathogens may trigger the production of disease-specific autoantibodies and arthritis in susceptible individuals. It has been shown recently that RA is associated with exposure to some microorganism such as *Aggregatibacter actinomyce-temcomitans* (Aa) [1].

In this study, we sought to conduct a large-scale survey to explore potential infectious agents which might initiate RA and the clinical consequence of this disease.

## Methods

### Patients

Survey results were collected from 902 RA patients admitted to the Department of Rheumatology and Immunology, People’s Hospital, Peking University, between December 2015 and June 2016. All the studied patients fulfilled the American College of Rheumatology/European League Against Rheumatism Classification criteria for RA in 2010, and written informed consent was obtained.

The clinical data were recorded including tender and swollen 28-joint counts, general health on visual analog scales, erythrocyte sedimentation rate (ESR), Health Assessment Questionnaire (HAQ)，28-joint Disease Activity Score (DAS28) and the infectious agents one month before RA onset.

The questionnaire including age, sex, disease duration, age at symptom, smoking status, DAS28 using the ESR at enrolment and treatments (one DMARD, more than one DMARDs, DMARDs plus low-dose glucocorticoid and bDMARDs). Only premorbid infectious agents of the RA patients were carefully recorded in this study.

### Statistical analyses

Analysis of covariance and multivariate logistic regression analysis were applied to compare the disease activity in patients with or without prior infections. T test or ANOVA was used to analyze the data. The categorical variables were compared with chi-squared test. Multinomial logistic analysis was used to find risk factor which perhaps affected the current disease activity in RA patients. Data was expressed as mean ± stand errors for continuous variables. The SPSS statistical package, version 23.0 was used for all statistical analyze, and p value less than 0.05 was considered statistically significant.

## Results

### 1. Prevalence of infections in RA

Within one month prior to RA onset, 37.03% (334/902) patients experienced infections, and the most frequent sites were respiratory (16.08%), intestinal (11.09%) and urinary (9.87%), respectively (Table 1).

**Table 1.** The type of premorbid infections in RA patients

| Infection types | Cases | Percentage (%) |
| --- | --- | --- |
| No infection | 568 | 62.97 |
| Respiratory | 145 | 16.08 |
| Upper | 120 | 13.30 |
| Lower | 25 | 2.77 |
| Intestinal | 100 | 11.09 |
| Urinary | 89 | 9.87 |

### 2. Patients in severe disease status showed high prevalence of infections

Four-hundred and ninety out of 902 RA patients with complete clinical data were analyzed in this study. These patients were divided into two groups based on DAS28 (DAS28<3.2 as group 1; DAS28≥3.2 as group 2). Compared with patients in group 2, patients in group 1 showed high prevalence of non-premorbid infection (X^2^=18.193，P=0.000) (Table 2). Notably, patients with high disease activity suffered more pre-exposure of respiratory, intestinal and urinary infections (P=0.000, P=0.000, P=0.023; respectively) (Table 2). Besides, higher ESR and CRP were observed in patients with higher DASD28 scores (Table 3).

**Table 2.** Prevalence of infection in RA patients with different disease activity

| Infection types | Group 1 (n=244) | Group 2 (n=246) | $\chi^2$ | P value |
| --- | --- | --- | --- | --- |
|  | DAS28<3.2<br>(n, %) | DAS28≥3.2<br>(n, %) |  |  |
| No infection | 201(82.4) | 161(65.4) | 18.193 | 0.000 |
| Respiratory | 20(8.2) | 39(15.9) | 30.384 | 0.000 |
| Intestinal | 10(4.1) | 17(16.9) | 125.390 | 0.000 |
| Urinary | 13(5.3) | 29(11.8) | 7.518 | 0.023 |

**Table 3.** Clinical characteristics and demographics of RA patients

| Characteristic |  | Group 1<br>(DAS28<3.2)<br>(n=244) | Group 2<br>(DAS28≥3.2)<br>(n=294) | Statistic | P value |
| --- | --- | --- | --- | --- | --- |
| Male<sup>c</sup>, n ( % ) |  | 50 (20.5) | 50 (20.3) | 4.601 | 0.100 |
| Age<sup>a</sup> ( years ) |  | 54±14 | 55±13 | -0.281 | 0.779 |
| Disease duration<sup>b</sup> ( years ) |  | 3 ( 2, 5 ) | 8 ( 3, 22.5 ) | -0.953 | 0.340 |
| Age at diagnosis<sup>b</sup> ( years ) |  | 44±14 | 45±15 | -0.781 | 0.435 |
| ESR<sup>b</sup> ( mm/H ) |  | 11 ( 7, 18 ) | 33 ( 19, 55 ) | -13.337 | 0.000 |
| CRP<sup>b</sup> ( mg/L ) |  | 2.68 (1.44, 4.87) | 9.58 (3.31, 23.19) | -10.531 | 0.000 |
| Anti-CCP negative<sup>c</sup>, n ( % ) |  | 37<br>(37/223, 16.6%) | 36<br>(36/154, 23.4%) | 2.686 | 0.101 |
| Anti-CCP antibody<sup>b</sup> ( U/L ) |  | 167.53<br>(57.2, 224.14) | 165<br>(38.24, 225.21) | -0.419 | 0.675 |
| HAQ<sup>b</sup> |  | 1 ( 0, 3 ) | 5 ( 1, 12 ) | -10.013 | 0.000 |
| Smoking Status<sup>c</sup> | Never smokers | 142<br>(142/209, 67.9) | 157<br>(157/215, 73.0) | 1.350 | 0.509 |
|  | Passive smokers | 23<br>(23/209, 11.0) | 19<br>(19/215, 8.8) |  |  |
|  | Active smokers | 44<br>(44/209, 21.1) | 39<br>(39/215, 18.1) |  |  |
| Current Treatment<sup>c</sup> | One DMARD | 56<br>(56/222, 25.2) | 38<br>(38/203, 18.7) | 15.904 | 0.001 |
|  | More than one DMARD | 138<br>(138/222, 62.2) | 109<br>(109/203, 53.7) |  |  |
|  | DMARDs + glucocorticoid | 14<br>(14/222, 6.3) | 23<br>(23/203, 11.3) |  |  |
|  | bDMARDs | 14<br>(14/222, 6.3) | 33<br>(33/203, 16.3) |  |  |

( ESR, erythrocyte sedimentation rate; CRP, C-reactive protein; anti-CCP, anti-citrullinated peptide antibodies; a, Data is described as mean ± SD, analysis with t-test; b, Data are reported as median with top and bottom quartile, nonparametric test is used for analysis; c, Chi-squared testis used. )

### 3. Disease Activity was associated with premorbid infection

In our study, patients showed higher DAS28 in urinary (P=0.000) and respiratory (P=0.001) infection groups (Table 4) before adjusting confounding factors such as the different therapies, age and smoking status which can affect disease activity.

**Table 4.** Differences of DAS28 between infectious groups and no infection group

|  | No-infection | Urinary | Respiratory | Intestinal |
| --- | --- | --- | --- | --- |
| Before adjusted | 3.25±0.07 | 3.97±0.19<sup>△</sup> | 3.78±0.17<sup>△</sup> | 3.56±0.26 |
| Adjusting for confounding factors |  |  |  |  |
| Therapy<sup>a</sup> | 3.20±0.07 | 3.91±0.20<sup>△</sup> | — | — |
| Age<sup>a</sup> | 3.26±1.41 | — | 3.80±1.450<sup>*</sup> | 3.58±1.50 |
| Smoking<sup>a</sup> | 3.24±1.38 | 3.95±1.56<sup>*</sup> | 3.82±1.50<sup>*</sup> | 3.62±1.51 |

Analysis of covariance was applied for adjusting confounding factors; a: Adjusted for therapy; b: Adjusted for age; c: Adjusted for smoking; —: Cannot be adjusted because of having an interaction effect compared with no-infection; Analysis of covariance was used between no-infection group and other infectious groups, \* : P<0.05 <sup>△</sup>: P<0.001.

One hundred and forty-five RA patients experienced respiratory tract infections one month prior to onset of the disease. Among these patients, 13.30% (120/902) patients showed upper respiratory tract infection while 2.77% (25/902) patients with lower respiratory tract infection. The number of tender and swollen joints (Fig 1A and B), HAQ scores (Fig 1D) and DAS 28 (Fig 1E) were higher in patients who had the respiratory tract infection compared with patients who had no infection before RA occurred. Furthermore, DAS28 was higher in respiratory infection group after adjusting for the age (P=0.002) and smoking (P=0.002) (Table 4)

**Fig 1.**
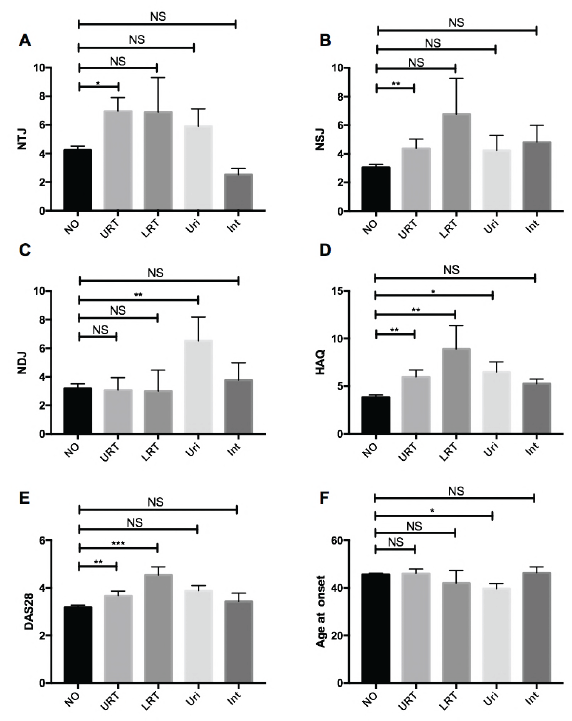
**Associations of disease activity and sites of infection**. These patients were categorical into subgroups including no infection (n=568), upper respiratory tract infection (n=120), low respiratory tract infection (n=25), urinary infection (n=89) or intestinal infection (n=100). Comparisons between groups were performed using the t-test or nonparametric test. The numbers of tender (A) and swollen (B) joints, HAQ (D) and DAS28 (E) were higher among respiratory tract infection, and number of deformity joints (C) was higher in urinary infection group. (F) Age at onset was younger in urinary infection group than other infection groups. (NO, no infection; URT, upper respiratory tract; LRT, lower respiratory tract; Uri, Urinary; Int, Intestinal)(*:P<0.05; **:P<0.01; ***:P<0.001)

There were 89 patients with urinary infection who developed RA in one month before disease initiation. More deformed joints (Fig 1C) were found in patients who had premorbid urinary infection. The age at onset was younger in patients who had urinary infection. (Fig 1F) DAS28 was still higher in urinary infection group after adjusting for the therapy type (P=0.000) and smoking (P=0.002) group (Table 4).

Intestinal infection occurred in 100 patients who developed RA. No difference was observed in these patients compared to patients with no infection. (Fig 1A-F) After adjusting age and smoking, DAS28 didn’t show significant difference between intestinal infection group and no infection group (Table 4).

### 4. Potential risk factors for high disease activity

The multinomial logistic regression was trained for predicting the disease activity with the factors which showed statistical significance in single-factor analysis (Supplementary Table 2). These model parameters were for the low, moderate and high levels of disease activity, measured relative to the remission level (reference outcome). High disease activity risk was increased in patients who had urinary infection (OR=3.813,95%CI=1.717-12.418) (Figure 2), and upper respiratory infection (OR=2.475, 95%CI= 0.971-6.312) (Figure 2).

**Figure 2.**
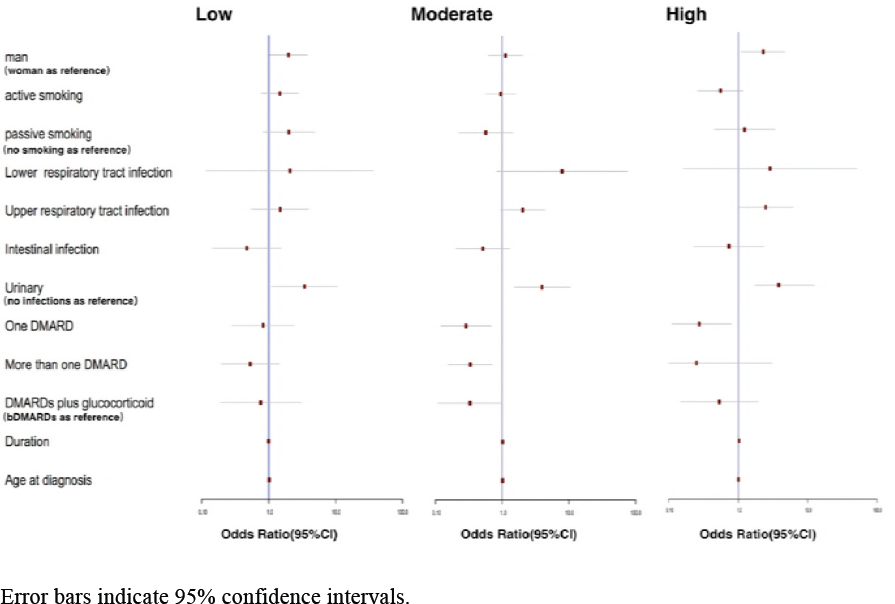
Multinomial Logistic regression for the potential risk factors for high disease activity. (Remission as reference)

## Discussion

There is increasing awareness that mucosal surfaces, including the gut and lungs, was sites of disease initiation in RA [8]. Recent studies showed that infectious agents including virus and bacteria infection had been associated with several kinds of autoimmune disease [7,10-12]. For instance, upper respiratory tract and other infections are well-known risk factors for multiple sclerosis [13]. However, it was not clearly whether infectious agents play the causative role in the onset or outcome of autoimmune disease, this is mainly due to the lack of strictly perspective epidemiological study. And even in animal models, these relationships are complex and depend on the timing of exposure, antigen type and genetic background [14]. In our study, the age of disease onset was younger in patients who had urinary tract infection, which perhaps indicates that RA occurred earlier in patients with this pre-exposure infection and later in the other patients.

It has been certified that many virus can play a role in the production of auto-antibodies such as anti-cyclic citrullinated peptide [15]. Infections are known to cause or enhance autoimmunity through expansion of auto-reactive T-cell clones by molecular mimicry and enhanced antigen presentation [14]. The patients with infection events during the disease duration could have advanced RA status [16]. To our knowledge, there was no study to prove the relationship between the premorbid infection history and onset or outcomes of RA in large populations. Here, we made the first report that analyzed this relationship in RA patients from outpatient of department of rheumatology and immunology in People’s Hospital, Peking University.

There were many factors reflected the disease activity in RA, such as the number of tender or swollen joints, ESR, CRP and so on. Patients with respiratory tract infection had higher DAS28 and more swollen/tender joints. This probably because of respiratory tract infection was mainly caused by viruses. Acute viral infection in adults have long been suggested to induce transient autoimmune responses, including generation of autoantibody [7]. As reported in a recent study, Arleevskaya et al found that higher percentages of first-degree healthy relatives (HR) than health control (HC) had upper respiratory and urinary tract infections. During10-year follow-up, 26 out of 251 (10.36%) HR subjects developed to RA, while no RA was found in HC group [4]. In our study, we found that 9.87% (89/902) patients had pre-exposure of urinary tract infection and 13.30% (120/902) patients with upper respiratory infection. Besides, the patients with urinary infection were more likely to stay in disease activity stage and have more deformity joints. Moreover, the patients with respiratory infection had higher disease activity compared with no infection patients.

In fact, it is impossible to make a causal link between a specific pathogen and the disease. Our study has several limitations. First, because the study was done in a retrospective manner, the patients who had no complete clinical data were excluded from this study. Second, the number of the studied patients was not large enough to see the statistical difference in clinical features and odds ratio in lower respiratory tract infection subgroup patients. It may be due to this study group with very few patients. Third, our studied patients may have selection bias because it was performed in a single university hospital. In order to determine the impact of premorbid infectious agents for RA outcome, the disease activity at RA onset and radiographic joint damage should be followed up in a larger prospective study.

## Acknowledgements

No.

## Reference

[1] Konig MF, Abusleme L, Reinholdt J, Palmer RJ, Teles RP, Sampson K, et al. Aggregatibacter actinomycetemcomitans-induced hypercitrullination linksperiodontal infection to autoimmunity in rheumatoid arthritis. SicTransl Med, 2016; 8(369):369ra176.

[2] Gray S. Firestein, Ralph C. Budd, Sherine E. Gabriel, et al. Kelley’s Textbook of Rheumatology. Singapore: Elsevier PteLtd, 2015:1139–1142.

[3] Arleevskaya MI, Kravtsova OA, Lemerle J, Renaudineau Y, Tsibulkin AP. How Rheumatoid Arthritis Can Result from Provocation of the Immune System by Microorganisms and Virus. Front Microbiol. 2016; 17(7):1296.

[4] Arleevskaya MI, Gabdoulkhakova AG, Filina YV, Miftakhova RR, Bredberg A, Tsybulkin AP. A transient peak of infections during onset of rheumatoid arthritis: a 10-year prospective cohort study. BMJ Open. 2014, 4(8):e005254.

[5] Leirisalo-Repo M. Early arthritis and infection. Curr Opin Rheumatol. 2005; 17: 433–439.

[6] Benedek TG. The history of bacteriologic concepts of rheumatic fever and rheumatoid arthritis. Semin Arthritis Rheum. 2006; 36(2):109–23.

[7] Ori Barzilai, Maya Ram, Yehuda Shoenfeld. Viral infection can induce the production of autoantibodies. Curr Opin Rheumatol. 2007; 19(6):636–43.

[8] Choe JY, Crain B, Wu SR, Corr M. Interleukin 1 receptor dependence of serum transferred arthritis can be circumvented by toll-like receptor 4 signaling. J Exp Med, 2003; 197:537.

[9] Liu Y, Mu R, Gao YP, Dong J, Zhu L, Ma Y, et al. A cytomegalovirus peptide-specific antibody alters natural killer cell homeostasis and is shared in several autoimmune diseases. Cell Host Microbe. 2016; 19(3):400–8.

[10] Brusca SB, Abramson SB, Scher JU. Microbiome and mucosal inflammation as extra-articular triggers for rheumatoid arthritis and autoimmunity. Curr Opin Rheumatol. 2014; 26:101–107.

[11] Haleniusm A, Henge H. Human cytomegalovirus and autoimmune disease. Biomed Res Int. 2014:472978. doi: 10.1155/2014/472978. Epub 2014 Apr 29

[12] IgoeA, Scofield RH. Autoimmunity and infection in Sjögren’s syndrome. Curr Opin Rheumatol. 2013; 25(4):480–7.

[13] Bulijevac D, Flach HZ, Hop WC, Hijdra D, Laman JD, Savelkoul HF, et al. Prospective study on the relationship between infections and multiple sclerosis exacerbations. Brain. 2002; 125(pt 5):952–60.

[14] MünzC, Lünemann JD, Getts MT, Miller SD. Antiviral immune responses: triggers of or triggered by autoimmunity? Nat Rev Immunol. 2009; 9(4): 246–258.

[15] Costenbader KH, Karlson EW. Epstein-Barr virus and rheumatoid arthritis: is there a link? Arthritis Res Ther. 2006;8:204

[16] Iguchi-Hashimoto M, Hashimoto M, Fujii T, Hamaguchi M, Furu M, Ishikawa M, et al. The association between serious infection and disease outcome in patients with rheumatoid arthritis. Clin Rheumatol. 2016; 35(1):213–8.

